# Uniformly re-analysed public omics data and graded meta-analyses of exercise responses in humans and rodents

**DOI:** 10.64898/2026.11.06.764830

**Authors:** Chaoyang Li, Qingqiu Su

## Abstract

Hundreds of omics studies have profiled molecular responses to exercise, but they differ in species, tissue, protocol, sampling time and analysis pipeline, so their results are hard to combine. This dataset places the public evidence on a common scale. We screened GEO, PRIDE and Metabolomics Workbench for exercise studies in humans, mice and rats, defined exercise–control contrasts from sample-level metadata and uniformly re-analysed 1,037 contrasts from 448 independent study clusters, covering 20 tissue classes and seven molecular data types. For every contrast, the dataset provides moderated standardized effect sizes for each gene, protein, miRNA or metabolite and, for expression data, pathways. For 88 analysis units defined by tissue, exercise paradigm and molecular layer, it provides random-effects meta-analyses of every feature measured in at least three independent studies, with heterogeneity statistics, evidence classes assigned by pre-specified criteria and replication tests in two held-out training atlases. Curation records, code and a static query website are included. The data support look-up of molecules, new meta-analyses and planning and benchmarking of exercise studies.

## Background & Summary

Regular physical activity prevents or attenuates many chronic diseases^1^ through molecular adaptations in many organs^2,3^. Over the past two decades, transcriptomic, epigenomic, proteomic and metabolomic profiling has been applied to exercised humans and rodents in hundreds of studies, and most of the resulting data have been deposited in public repositories. Individual studies are typically small — the median transcriptomic contrast in this dataset has five samples in its smaller group — and they differ in species, exercise modality and dose, sampling time, tissue, health context and analysis pipeline. Results of small and heterogeneous studies are often not reproduced^4,5^, and published lists of exercise-responsive molecules cannot simply be compared, because each list reflects its own design, platform and processing.

Several resources address parts of this problem. Meta-analyses of human skeletal muscle and blood transcriptomes^6,7^ and multi-omic analyses of large human cohorts^8,9^ provide pooled estimates for human muscle; web portals collect exercise datasets for browsing and re-analysis^10–12^; and the Molecular Transducers of Physical Activity Consortium (MoTrPAC)^13,14^ and a recent 22-tissue mouse atlas^15^ have generated deep multi-omic maps of standardized training protocols. The dataset described here complements these resources. It covers every eligible exercise series that we identified in three repositories, across tissues, species and molecular layers; it records how each exercise–control comparison was defined from the sample-level metadata; it re-analyses the primary data with uniform pipelines; and it pairs each pooled estimate with measures of between-study consistency, a pre-specified evidence class and, where possible, tests in independent data.

Figure 1 summarizes how the dataset was generated. We searched GEO, PRIDE and Metabolomics Workbench on 24 October 2026; screened 1,478 GEO series, 254 PRIDE projects and 215 Metabolomics Workbench studies; and defined 1,309 exercise–control contrasts from GEO sample records (1,088 for discovery and 221 from two held-out atlases), together with 29 proteomic and 39 metabolomic contrasts. The sample selection of every contrast was verified programmatically against the repository metadata. We re-analysed the primary data of 1,037 discovery contrasts from 448 independent study clusters, computed moderated standardized mean differences for every feature and, for expression data, every pathway, meta-analysed each feature measured in at least three independent clusters within units defined by tissue, exercise paradigm, modality and molecular layer, and assigned it to one of six evidence classes using criteria adapted from umbrella reviews^16,17^ and fixed before any effect size was computed. The classes were then tested against the held-out MoTrPAC rat atlas^14^ and mouse atlas of Maier et al.^15^ and by repeated split-half replication.

**Fig. 1.**
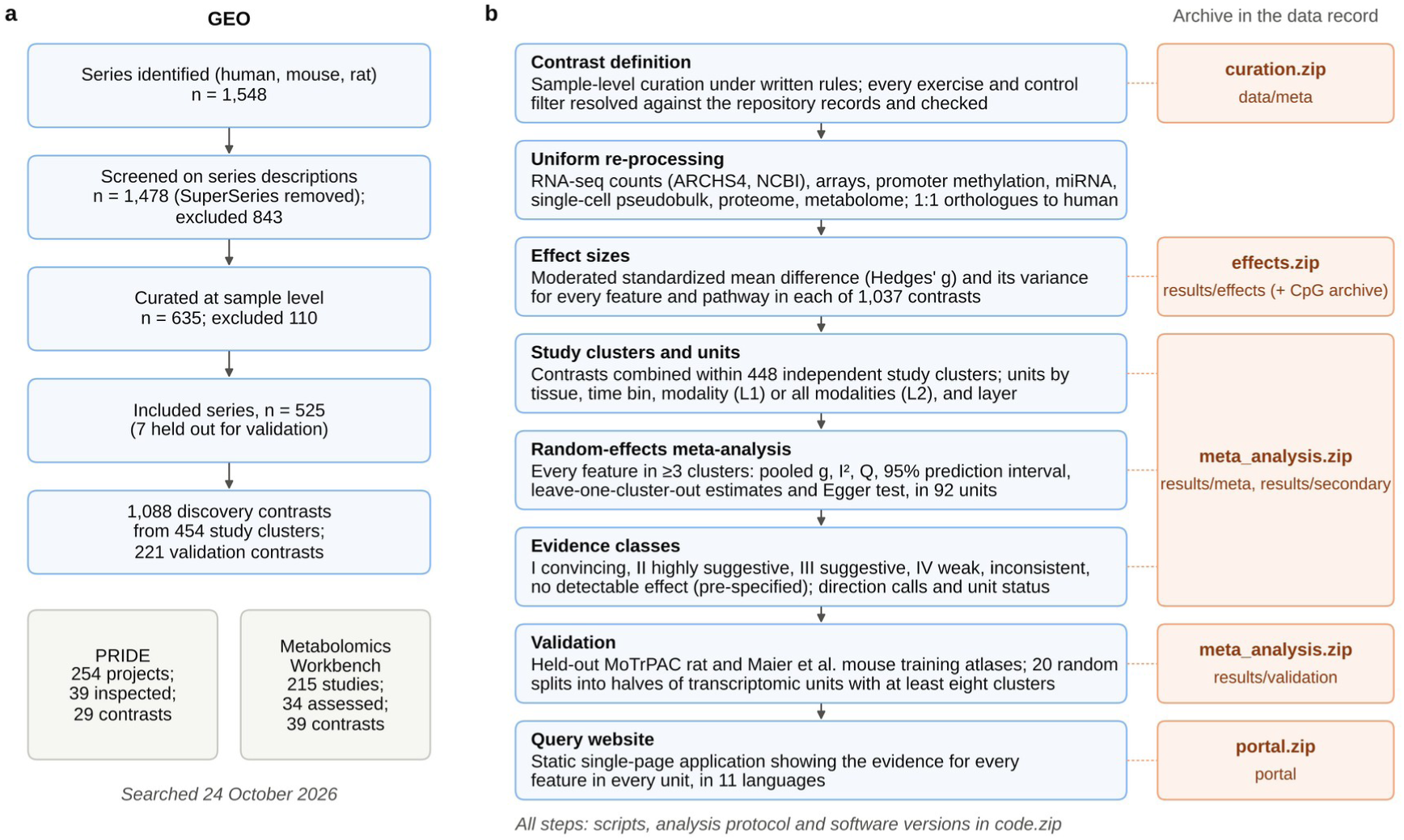
Generation of the dataset. **a** Identification, screening and inclusion of GEO series, with the resulting discovery and validation contrasts and study clusters, and of PRIDE projects and Metabolomics Workbench studies. Series generated by MoTrPAC and by the mouse atlas of Maier et al. were held out for validation. **b** Processing steps from the definition of exercise–control contrasts to the query website (left; split-half validation was done in transcriptomic units with at least eight clusters) and the archive of the data record that holds the output of each step (right). L1, modality-specific units; L2, units pooling all exercise modalities; IZ, between-study heterogeneity; Q, Cochran’s heterogeneity statistic.

The data can be reused in several ways. The pooled effect, heterogeneity, prediction interval, evidence class and external replication of any gene, protein, miRNA or metabolite can be looked up for each tissue and exercise paradigm, so that a candidate molecule can be checked against cross-study evidence before it is followed up. The per-contrast effect sizes and their harmonized annotations allow new syntheses — for example by sex, age group, health context or platform, or after adding new studies — without re-processing the primary data. Between-study heterogeneity and prediction intervals indicate the range of effects that a new study can expect and can inform sample-size calculations. The curation tables record why each series was excluded or could not be re-analysed and which combinations of tissue, paradigm and molecular layer have been studied, which shows where new data are needed. Finally, features with high evidence classes, together with their replication status, provide reference sets for benchmarking computational methods and for comparing new exercise studies with previous work. These data have not been described in a previous publication.

## Methods

### Overview and analysis protocol

The dataset was generated by a data-driven synthesis that combines the search and screening steps of a systematic review with uniform re-analysis of primary data. An internal analysis protocol, which was not registered, specified the eligibility criteria, the unit of analysis, the effect-size and meta-analytic models, the evidence classes and the discovery–validation split; implementation details (unit levels, tissue harmonization, timepoint binning and the definition of independence) were added before any effect size was computed. The protocol and its log of deviations are part of the data record (protocol/PROTOCOL.md in the code archive); implementation choices that the protocol did not cover, such as the source of RNA-seq counts and the expression filters, are described below. Search and screening are reported following PRISMA 2020 where applicable^18^ (Fig. 1a).

### Data sources and search

NCBI GEO^19^ was searched through the E-utilities on 24 October 2026 for series (entry type gse) from *Homo sapiens*, *Mus musculus* or *Rattus norvegicus* that matched, in any field, at least one of 28 terms: exercise, “physical activity”, treadmill, “wheel running”, “voluntary running”, “running wheel”, “endurance training”, “resistance training”, “resistance exercise”, “strength training”, “interval training”, HIIT, “swimming training”, “swim training”, “swimming exercise”, “exercise training”, “physical training”, “aerobic training”, “aerobic exercise”, athletes, marathon, “physically active”, exercised, “trained rats”, “trained mice”, “ladder climbing”, “cycling exercise” and “exercise intervention”. Series- and sample-level metadata (SOFT ‘brief’ records) were retrieved for all 1,548 hits. The 70 SuperSeries among them were removed, so that their SubSeries were screened only if the search had returned them; 54 of the 197 SubSeries of these SuperSeries had not been returned and were not screened. PRIDE^20^ projects were identified through the PRIDE Archive API with 16 keywords (exercise, endurance training, resistance training, treadmill, wheel running, physical activity, high-intensity interval, HIIT, running mice, exercise training, trained rats, swimming training, acute exercise, sprint, cycling exercise and athletes), and Metabolomics Workbench^21^ studies through its REST interface by title search with 13 keywords (exercise, training, running, treadmill, physical activity, endurance, athlete, HIIT, resistance, marathon, cycling, sprint and wheel). The search code, which contains the query strings, and all returned records are included in the data record.

### Eligibility

We included in vivo exercise in humans, mice or rats with a within-study comparator, in three designs: acute bouts (post-versus pre-exercise in the same individuals, or exercised versus rested animals); training interventions (post-versus pre-training, or trained versus sedentary animals); and habitually trained versus untrained people (cross-sectional, analysed separately). Exercise in disease, ageing or diet models was eligible when exercised and sedentary animals shared the same background; in studies of engineered genotypes, only wild-type arms were used. We excluded in vitro and ex vivo stimulation models, pharmacological exercise mimetics, electrical stimulation, disuse-only designs, offspring designs, tumour tissue, contrasts in which exercise was confounded with another factor, and groups with fewer than two samples. Bed rest with an exercise countermeasure, compared with bed rest alone, was eligible and coded as health context ‘bed rest’. Exclusions were coded with 11 pre-defined reasons, which are recorded for every series.

### Screening and contrast definition

Screening was performed in two stages by a large language model working under explicit written rules, which are included verbatim in the curation archive. The sample selection of every contrast was verified programmatically (see below); the other annotations were not independently audited. Stage 1 classified each series from its title, summary, overall design and sample sources. Stage 2 used compact tables of all sample-level characteristics and, for every eligible contrast, recorded the species, tissue (free text and a harmonized major tissue class), molecular layer, paradigm, modality (endurance, END; voluntary wheel running, VWR; resistance, RES; high-intensity interval, HIIT; combined, CON; other, OTH), time after the end of exercise, training duration, sex, age group, health context and design (paired or independent groups), and the exercise and control sample filters, expressed as pairs of a metadata field and a regular expression. Every filter was resolved programmatically against the GEO sample records, and a contrast was retained only if both groups had at least two samples, no sample fell in both groups and the numbers of selected samples matched the group sizes expected by the curator. For paired designs, subject identifiers were extracted with a capture expression, and contrasts with fewer than three matched pairs were analysed as independent groups. Series generated by MoTrPAC or by Maier et al. were flagged as validation sources and removed from discovery. The same rules were applied to PRIDE result tables (mapping of table columns to groups) and to Metabolomics Workbench studies (mapping of sample factors to groups); the curator’s group definitions and expected group sizes for these contrasts are recorded in the PRIDE and Metabolomics Workbench curation files.

Acute contrasts were assigned to time bins by the time from the end of exercise to sampling: A0, ≤1 h; A1, >1–12 h; A2, >12–72 h. Training contrasts formed bin C and cross-sectional comparisons bin X; acute contrasts with unknown timing (Aunk) or sampled more than 72 h after exercise (Alate) were catalogued but not assigned to analysis units.

Independence was defined at the level of study clusters, which group GEO series that share a PubMed identifier, a curated cohort annotation, or a first contributor within ±1 submission year. Each PRIDE project and Metabolomics Workbench study formed its own cluster and was not merged with GEO clusters; two PRIDE projects from the same study, whose cohorts may overlap (PXD022262 and PXD027406), were therefore treated as separate clusters.

### Uniform re-processing

For human and mouse bulk RNA-seq, we used uniformly processed gene-level counts from ARCHS4 (version 2, July 2026 release)^22^, read remotely by sample column; contrasts with any sample absent from ARCHS4 used NCBI GEO-generated counts, and the remaining contrasts, including all rat contrasts, used submitter count tables, whose columns were matched to samples automatically or, where this failed, by curator-defined maps. Counts were normalized by the trimmed mean of M-values (TMM)^23^ within each contrast and transformed to log_2_ counts per million; genes with ≥1 count per million in at least as many samples as the smaller group were retained. When only normalized values (TPM or FPKM) were available, we used log_2_(*x* + 1) and removed the 30% of genes with the lowest mean expression.

For expression arrays, we used submitter-normalized series-matrix values, log_2_-transformed linear-scale data and quantile-normalized all arrays within each contrast. Probes were mapped to Entrez Gene through the GEO platform annotation or, where only accessions were provided, through gene symbols parsed from RefSeq descriptions or NCBI gene2refseq; multi-mapping probes were removed, the probe with the highest mean intensity was kept for each gene, and the 20% of genes with the lowest intensity were removed. For platforms with custom probe-set definitions whose identifiers encode genes (Entrez or Ensembl gene identifiers, or Ensembl transcript identifiers mapped through NCBI gene2ensembl) or are gene symbols, identifiers were mapped directly. Submitter tables that required more than a single column- to-sample map — contrasts split across files or worksheets, transcript-level tables (summed to genes), non-standard headers or paired designs with samples absent from some files — were mapped per contrast by the curator and verified programmatically. Transcriptomic contrasts with fewer than 3,000 genes after filtering were not used.

For Illumina 450K and EPIC DNA methylation arrays, series-matrix β values were read in probe blocks, restricted to autosomal CpG probes with <20% missing values and converted to M-values; gene-promoter methylation was the mean M-value of probes annotated to TSS1500, TSS200, the 5′ untranslated region or the first exon, requiring at least two probes (EPIC probes shared with the 450K array were annotated with the 450K manifest). When series matrices contained no data, processed β-value tables from the supplementary files were used after curator-defined column-to-sample mapping; EPIC v2 probes were collapsed onto their 450K/EPIC identifiers; one transposed table (samples as rows) was aggregated to promoters sample by sample; and one PRE/POST array pair that was byte-identical in the submitted table was excluded. Mouse methylation BeadChips, whose platform tables carry probe coordinates but no gene annotation, were assigned to promoters (transcription start site −1,500 to +500 bp) using NCBI RefSeq transcription start sites for the mm10 assembly. For sequencing-based methylation (RRBS and WGBS), per-CpG or per-region methylation from processed tables or Bismark coverage files (CpGs with ≥5 reads) was converted to M-values and averaged over the same promoter window (≥2 CpGs), using RefSeq transcription start sites for the genome build stated by the submitters (hg19, hg38, mm10 or mm39; one mm9 data set was used through its own RefSeq promoter assignment); MeDIP-seq enrichment data were not used. Array- and sequencing-based methylation were treated as separate layers.

miRNA arrays were mapped to mature miRNA names through platform annotations, small-RNA-seq tables were parsed from supplementary files (with curator-defined column-to-sample maps where automatic parsing failed), and names were harmonized by removing species prefixes; tables keyed by precursor gene symbols (for example, MIR21) or Ensembl gene identifiers were converted to precursor names. For single-cell and single-nucleus RNA-seq, per-sample 10x count matrices were summed over all cells (pseudobulk) and analysed as count data in a separate single-cell layer; data sets deposited only as R objects could not be read. For proteomes, we used protein-level intensity tables (MaxQuant, DIA-NN, Spectronaut or vendor reports, including tables inside zip or 7z archives, identified in two passes over all exercise-related PRIDE projects, with sample-design files used to resolve groups), log_2_-transformed and median-centred them, kept proteins quantified in ≥70% of the samples of each group and mapped them to genes through gene names or UniProt accessions. Named metabolites (RefMet nomenclature) were retrieved through the Metabolomics Workbench REST interface, log_2_-transformed, median-centred per sample and filtered in the same way; when a metabolite was measured on several analytical platforms within a study, the most precise estimate was kept.

Mouse and rat genes were mapped to human Entrez Gene identifiers using 1:1 orthologues from NCBI gene_orthologs. The reason why each eligible contrast could not be re-analysed was recorded in one of 13 categories.

### Effect sizes

For each contrast and feature, we computed a moderated standardized mean difference (Hedges’ *g*) in which gene-wise variances were shrunk towards a common prior by empirical Bayes moderation (limma squeezeVar^24^), following the moderated effect-size approach of ref. 25. For independent groups, *g* = *J*(*x^⎯^*_E_ – *x^⎯^* _C_)/*s*∼_pooled_, with *J* = 1 − 3/(4*df* − 1) and var(*g*) = 1/*n*_E_ + 1/*n*_C_ + *g*^2^/[2(*n*_E_ + *n*_C_)]. For paired designs^26^, the mean difference was standardized by the average within-condition standard deviation (*d*_av_), which places paired and independent estimates on the same metric, with var(*g*) = (1/*n* + *g*^2^/2*n*) × *s*^∼^_D_^2^/*s*^∼^_av_^2^, where the variance ratio equals 2(1 − *r*) and was bounded to [0.1, 2]. In both cases, a positive *g* denotes a higher value after exercise, or in trained than in untrained or control samples (for DNA methylation, higher promoter methylation). Pathway activity scores for MSigDB Hallmark (version 2024.1)^27^ and Reactome^28^ gene sets of 15–300 genes were computed per sample as the mean of within-contrast *z*-scored member genes (requiring at least ten measured members and one third of the set), and pathway effect sizes were computed in the same way as gene effect sizes.

### Analysis units and meta-analysis

Analysis units were defined at two levels: level 1 (L1, modality-specific: major tissue class × time bin × modality × molecular layer) and level 2 (L2, exercise-general: all modalities pooled). Expression arrays and RNA-seq formed one transcriptome layer. Within a unit, contrasts from the same study cluster were combined into a single estimate: contrasts sharing samples (for example, several time points or exercise arms against one control group) were averaged, treating them as fully dependent, and contrasts on independent samples were pooled by inverse-variance weighting. Units with at least three independent clusters were analysed.

Each feature measured in at least three study clusters within a unit was meta-analysed with a random-effects model, with the between-study variance τ^2^ estimated by restricted maximum likelihood (fixed-point iteration)^29^. We recorded the pooled estimate and its standard error, the Wald *z* test, Cochran’s *Q* and *I*^2^ (ref. 30), the 95% prediction interval (*t* distribution with *k* − 2 degrees of freedom)^31^, the numbers of clusters with positive, negative and nominally significant effects, leave-one-cluster-out estimates and, for *k* ≥ 10, Egger’s regression test^32^. False discovery rates (Benjamini–Hochberg) were computed within each unit.

### Evidence classes

Evidence classes were pre-specified by adapting the credibility criteria of umbrella reviews^16,17^ to genome-wide testing (Table 1). Each feature was assigned the highest class whose criteria it met: class I (convincing), class II (highly suggestive), class III (suggestive) or class IV (weak). Features with *k* ≥ 3 and *P* ≥ 0.05 were classed as inconsistent if at least one cluster was nominally significant in each direction or *I*^2^ ≥ 75%, and otherwise as showing no detectable effect; features measured in fewer than three clusters were insufficient. Classes rate how credible a change is, irrespective of its direction, which is given by the sign of the pooled *g*.

**Table 1.** Criteria for the evidence classes.

| Class | Clusters (*k*) | Significance | Further criteria |
|---|---|---|---|
| I, convincing | ≥5 | FDR < 10<sup>−3</sup> | *I*<sup>2</sup> < 50%; 95% prediction interval excludes zero; ≥80% of clusters concordant in sign; all leave-one-cluster-out estimates nominally significant in the same direction; Egger *P* > 0.10 when *k* ≥ 10 |
| II, highly suggestive | ≥3 | FDR < 0.01 | ≥70% of clusters concordant in sign; ≥80% of leave-one-cluster-out estimates nominally significant in the same direction |
| III, suggestive | ≥3 | FDR < 0.05 | – |
| IV, weak | ≥3 | *P* < 0.05 | – |
| Inconsistent | ≥3 | *P* ≥ 0.05 | At least one cluster nominally significant in each direction, or *I*<sup>2</sup> ≥ 75% |
| No detectable effect | ≥3 | *P* ≥ 0.05 | Not inconsistent |
| Insufficient | <3 | – | – |

Each feature receives the highest class whose criteria it meets. FDR, Benjamini–Hochberg false discovery rate within the analysis unit; P, P value of the pooled random-effects estimate.

Each feature in each unit was also given a direction call: up or down for classes I–III and possibly up or possibly down for class IV (sign of the pooled *g*); conflicting for the inconsistent class; and, for features without a detectable effect, no change when the whole 95% confidence interval lay within ±0.2 (a negligible standardized difference) or no detectable change otherwise. Each combination of tissue, time bin and layer was also assigned a summary status. A unit was established if at least ten features were graded I–III and the grades were confirmed in held-out data — a Cochran–Armitage trend *z* > 1.96 in every available external atlas or, in split halves, class I–II features replicating at least twice as often as features without a detectable effect, with at least ten class I–II features per half — unless the external trend was ≤0 in every atlas. Otherwise, a unit was contested if the calibrated definition of inconsistency (see below) applied to at least 25% of features without a significant pooled effect, tentative if at least ten features were graded I–III, and showing no consistent effect if none of these applied. Combinations with fewer than three re-analysed clusters, or without any feature measured in three, had too few studies. For layers other than the transcriptome, the acute time bins were combined and the most informative status was shown. The direction calls and summary statuses were defined after the primary analyses, as summaries for the query website.

### Validation

External validation used two held-out resources (Table 2). For MoTrPAC (rats; 8 weeks of progressive treadmill training versus sedentary controls)^14^, we used the consortium’s differential-analysis results (MotrpacRatTraining6moData^33^) for the transcriptome, proteome and metabolome, combined sex-specific log fold changes by fixed-effect pooling and pooled tissues within a major class (for example, gastrocnemius and vastus lateralis); proteins were mapped from RefSeq protein accessions to rat genes (NCBI gene2refseq) and then to human orthologues. For the mouse atlas of Maier et al. (8 weeks of voluntary wheel running)^15^, we re-analysed the deposited bulk RNA-seq data^34^ with the discovery pipeline and combined sexes and circadian phases by fixed-effect pooling. A discovery feature was considered replicated if the validation estimate had the same sign and *P* < 0.05. Replication across classes I, II, III, IV and no detectable effect was tested with the Cochran–Armitage trend test, and effect sizes were compared by Spearman correlation. Because the validation estimates pool sexes, circadian phases and several tissues of the same animals (for example, nine skeletal muscles in the mouse atlas) by fixed effect, their standard errors are underestimated; replication rates are therefore best compared between classes rather than read as absolute probabilities. For internal validation, the study clusters of each transcriptomic unit with at least eight clusters (32 units) were split at random into two halves 20 times; features were graded in one half (without the Egger criterion) and counted as replicated if the other half gave an estimate of the same sign with *P* < 0.05.

**Table 2.** Held-out resources used for validation.

| Resource | Species and design | Tissues and layers used | Data used |
|---|---|---|---|
| MoTrPAC endurance exercise training study<sup>14</sup> | Rats; 8 weeks of progressive treadmill training versus sedentary controls; both sexes | Tissues mapped to the discovery tissue classes and pooled within each class (for example, gastrocnemius and vastus lateralis); transcriptome, proteome, metabolome | Consortium differential-analysis results (MotrpacRatTraining6moData<sup>33</sup>) ; sex-specific log fold changes combined by fixed-effect pooling |
| Multi-omics atlas of chronic exercise, Maier et al.<sup>15</sup> | Mice; 8 weeks of voluntary wheel running versus sedentary controls; both sexes, sampled in two circadian phases | Tissues mapped to the discovery tissue classes and pooled within each class (for example, nine skeletal muscles); transcriptome | Deposited bulk RNA-seq data (GSE344069<sup>34</sup>) re-analysed with the discovery pipeline; sexes and phases combined by fixed-effect pooling |

GEO series of both resources (MoTrPAC, GSE242354, GSE242355 and GSE242357; Maier et al., GSE344068, GSE344069, GSE344070 and GSE347387) were excluded from discovery.

### Additional derived tables

The data record also contains tables from secondary analyses. Analyses specified in the protocol were sensitivity analyses (restriction to healthy contexts, to independent-group designs or to RNA-seq, and exclusion of the largest cluster), which were extended with human-only, rodent-only, array-only and paired-design analyses (protocol deviation D5) and run in 11 major transcriptomic units with ≥10 clusters chosen to cover the main tissues and paradigms; mixed-effects meta-regression of pathway effects on study-level moderators (human versus rodent, healthy versus disease context, RNA-seq versus arrays, inclusion of females); Spearman correlations of unit-level estimates between molecular layers; and counts of features graded I–II in several tissues for the same time bin. The following were added after inspection of the primary results and are labelled as post hoc in the data record: a calibrated definition of inconsistency, which additionally requires Cochran’s *Q P* < 0.05, and the share of features expected to meet the primary rule under homogeneity, 1 − 2(0.975)*^k^* + 0.95*^k^* averaged over the non-significant features of each unit; the minimal detectable pooled |*g*| (80% power, two-sided *P* = 10^−4^), approximated as 4.73 times the median standard error of pooled estimates; the genomic inflation factor λ of the contrasts in ten major transcriptomic units, defined as the median of *g*^2^/var(*g*) across genes divided by 0.4549, and sensitivity analyses excluding contrasts with λ > 2; the heterogeneity of 17 core clock genes and 12 glucocorticoid-receptor target genes relative to all other genes; gene type and orthology by class; re-analysis of endurance-trained muscle restricted to human or to rodent clusters and comparison with resistance training and with the held-out atlases; comparison of muscle training effects with the same genes in other tissues; look-up of 26 gene–unit combinations reported in earlier landmark studies; and the reporting completeness of the contrast annotations. The scripts that produce these tables are included in the code archive.

### Query website

A static single-page website was generated from the unit-level tables (code/39_portal_data.py and code/40_portal_i18n.py). For every feature measured in at least three independent studies in a unit, it shows the direction call, evidence class, pooled *g* with 95% confidence and prediction intervals, heterogeneity, contributing studies and external replication, together with the summary status of each unit, an evidence map and definitions of all terms, in 11 languages. The website has no server-side code and loads sharded JSON files from its data folder.

### Software and use of AI assistance

All analyses were implemented in Python 3.13 (NumPy, SciPy, pandas, h5py, pyreadr and matplotlib; versions are listed in requirements.txt in the code archive). Screening, sample-level annotation and table-to-sample mapping were performed by a large language model (Claude, Anthropic) under written rules, and every resulting sample filter and column map was verified programmatically against the source metadata. Analysis code was written with the same assistance and reviewed by the authors, who take full responsibility for the content.

### Ethics statement

All data analysed were publicly available and de-identified, and no new data were collected from humans or animals; ethical approval was therefore not required.

## Data Records

The data record is available at Zenodo^35^ (version 1.0, published 25 October 2026). Data are released under the Creative Commons Attribution 4.0 International licence (CC BY 4.0) and code under the MIT License. The record comprises a README file and six zip archives (Table 3; 766.8 MB in total), which unpack into a common folder structure (data/meta, results, code, protocol and portal). All tables are tab-separated text with a header row, gzip-compressed where the file name ends in .gz.

**Table 3.** Files in the data record.

| File | Size | Content |
|---|---|---|
| README.md | 5.0 kB | Overview of the record, file formats, licences and instructions for reproducing the analyses |
| ExGrade_v1.0_curation.zip | 4.1 MB | data/meta: search results and repository metadata, screening decisions, contrast definitions, reasons for non-re-analysis, curator instructions and outputs, column-to-sample maps (153 files) |
| ExGrade_v1.0_effects.zip | 229.3 MB | results/effects: per-contrast effect sizes for every feature and pathway in seven molecular layers (2,284 files) |
| ExGrade_v1.0_effects_methylation_cpg.zip | 354.5 MB | results/effects/methylation/cpg: per-contrast CpG-level effect sizes for methylation arrays; intermediate files (41 files) |
| ExGrade_v1.0_meta_analysis.zip | 167.3 MB | results/meta: unit-level meta-analyses with evidence classes and unit summaries; results/validation: external and split-half validation; results/secondary: additional derived tables; results/numbers.json (466 files) |
| ExGrade_v1.0_portal.zip | 11.4 MB | portal: static query website (135 files) |
| ExGrade_v1.0_code.zip | 240.9 kB | code: analysis scripts; protocol/PROTOCOL.md: analysis protocol and deviations log; requirements.txt; LICENSE (67 files) |

### Identifiers

Genes are identified by human NCBI Entrez Gene identifiers. Mouse and rat genes were mapped to human identifiers through 1:1 orthologues, so genes without a 1:1 human orthologue are absent from rodent contrasts. Proteins are reported at the gene level with the same identifiers, miRNAs by lower-case names without species prefix (mature names such as mir-21-5p, or precursor names such as mir-21 where only precursor-level data were deposited), metabolites by lower-case RefMet names and pathways by their MSigDB Hallmark or Reactome names. A contrast is identified by its source accession (GEO series, PRIDE project or Metabolomics Workbench study) and a contrast number joined by two underscores (for example, GSE104079 c2), and a study cluster by ‘SC’ and four digits (PRIDE and Metabolomics Workbench clusters by ‘PX_’ or ‘MW_’ followed by the project or study accession). Analysis units are named layer|level|tissue|time bin|modality (for example, transcriptome|L2| skeletal_muscle|C|ALL), with ’ ’ replacing ‘|’ in file names.

### Curation archive (ExGrade_v1.0_curation.zip; 153 files in data/meta)

This archive contains: the search results and metadata, geo_series.tsv (1,548 series, with title, summary, overall design, contributors, PubMed identifiers and SuperSeries relations) and geo_samples.tsv (61,441 samples, with title, source, characteristics and description), and the PRIDE and Metabolomics Workbench candidate lists (pride_candidates.tsv, mwb_candidates.tsv); the screening decisions, screen1_all.tsv (stage 1 decision, exclusion code and provisional annotation for the 1,478 screened series) and stage2_series_decisions.tsv (stage 2 decision, exclusion code, validation flag and number of contrasts for 635 series); the contrast definitions, contrasts_resolved.tsv (one row for each of the 1,309 GEO contrasts; Table 4) and contrasts_proteome.tsv and contrasts_metabolome.tsv (corresponding fields for the 29 PRIDE and 39 Metabolomics Workbench contrasts, without the filter and expected-size fields; group members are listed as table column names or Metabolomics Workbench sample identifiers, and the curator’s group definitions and expected group sizes are in pride_curation.jsonl, pride_curation2.jsonl and mwb_curation.jsonl); unprocessed_reasons.tsv (category and detail for each of the 119 eligible GEO discovery contrasts that were not re-analysed); and the curator instructions, curator outputs and curator-defined column-to-sample maps for each stage (folders screen1, screen2, suppmap, methmap and methseqmap, and the pride_ and mwb_ files).

**Table 4.** Main fields of the contrast table and of the unit-level meta-analysis tables.

| Field | Description |
|---|---|
| **contrasts_resolved.tsv** | *One row per GEO contrast; contrasts_proteome.tsv and contrasts_metabolome.tsv have corresponding fields, without the filter and expected-size fields (see text)* |
| gse, cid | Source accession and contrast number; the contrast identifier is gse__cid |
| species | human, mouse or rat |
| tissue, tissue_detail, tissue_major | Tissue as described by the submitters, further detail, and harmonized major tissue class |
| layer_raw, layer, platform | Data type as curated, analysis layer, and GEO platform accession |
| paradigm, tbin | acute, chronic (training) or cross_sectional; time bin: A0 (≤1 h), A1 (>1–12 h), A2 (>12–72 h), Aunk (unknown), Alate (>72 h), C (training), X (cross-sectional) |
| timepoint_h, duration_wk | Hours from the end of exercise to sampling (acute contrasts; also recorded for 64 training contrasts); training duration in weeks |
| modality | END, VWR, RES, HIIT, CON or OTH |
| sex, age_group, health | male, female, both or unknown; young, adult, middle, older, aged or unknown; healthy or the disease, diet or other model context |
| design, pairs | paired or independent; matched (control, exercise) sample pairs for paired designs |
| exercise_filter, control_filter | Curator-defined sample filters: lists of (metadata field, regular expression) pairs that a sample must all match |
| gsm_ex, gsm_ctl | Samples of the two groups: GEO sample accessions selected by the filters (table column names for PRIDE, sample identifiers for Metabolomics Workbench) |
| n_ex_expected, n_ctl_expected, n_ex, n_ctl, n_pairs | Group sizes expected by the curator and resolved from the filters; number of matched pairs |
| pubmed, contributors, submission, cohort_hint | Fields used to define study clusters |
| study_cluster, validation_source | Independent study cluster (SC####; PX_ or MW_ plus accession for PRIDE and Metabolomics Workbench); True for series from the held-out atlases |
| status, dup_with | Resolution status (‘ok’ for all retained contrasts); other series sharing samples (none) |
| **<unit>__genes.tsv.gz** | *One row per feature in an analysis unit; <unit>__pathways.tsv.gz has the same fields for pathways* |
| gene | Feature identifier: Entrez Gene identifier, miRNA name or metabolite name |
| k, N_total | Number of independent study clusters and of samples (or pairs) contributing |
| mu, se, z, p, fdr | Pooled Hedges’ *g* (random effects), standard error, Wald *z*, *P* value and Benjamini–Hochberg false discovery rate within the unit |
| tau2, I2, Q, Qp | Between-study variance, *I*<sup>2</sup> (%), Cochran’s *Q* and its *P* value |
| pi_lo, pi_hi | Lower and upper limits of the 95% prediction interval |
| n_pos, n_neg, n_sig_pos, n_sig_neg | Numbers of clusters with positive or negative estimates, and with nominally significant (*P* < 0.05) positive or negative estimates |
| sign_agree | Share of clusters whose estimate has the sign of the pooled estimate |
| loo_same_frac, loo_sig_frac | Share of leave-one-cluster-out estimates with the same sign as the pooled estimate, and with the same sign and *P* < 0.05 |
| egger_p | *P* value of Egger’s regression test (*k* ≥ 10) |
| class | Evidence class: I, II, III, IV, Inconsistent, No effect or Insufficient |

### Effect-size archive (ExGrade_v1.0_effects.zip; 2,284 files)

The folders results/effects/<LAYER>/genes and results/effects/<LAYER>/pathways contain one gzip-compressed table per contrast, named <CONTRAST>.tsv.gz, for the seven layers transcriptome, miRNA, methylation (arrays), methylation_seq, singlecell, proteome and metabolome. Feature tables have the columns gene (feature identifier: Entrez Gene identifier, miRNA name or metabolite name), g (Hedges’ *g*), v (sampling variance of *g*), diff (difference in means, or mean within-pair difference, on the analysis scale: log_2_ expression or intensity, or M-value for methylation) and, for expression data, mean (average value on the analysis scale). Pathway tables have the columns pathway, g, v and diff; they contain values for transcriptome, single-cell and proteome contrasts and are empty for the other layers. Fifteen proteome files contain one row with an empty identifier (proteins that could not be mapped to a gene); these rows, and the corresponding row of the proteome meta-analysis, should be ignored. The transcriptome folders also contain the 104 re-analysed contrasts of the held-out mouse atlas (GSE344069, chronic wheel running, 88 contrasts; GSE344070, acute wheel running, 16 contrasts), which are flagged in the validation_source column of contrasts_resolved.tsv; only GSE344069 was used for validation. The metabolome folders contain one further file, ST003662 m1, left from an earlier processing run; the sample filters of this contrast did not resolve in the final run, so it is not part of contrasts_metabolome.tsv and is not used in any analysis.

#### CpG-level methylation archive (ExGrade_v1.0_effects_methylation_cpg.zip; 41 files)

results/effects/methylation/cpg contains CpG-level effect sizes (columns as above, keyed by array probe identifier) for the methylation-array contrasts that were processed from probe-level data. These are intermediate files; the promoter-level values used in the meta-analyses are in the effect-size archive.

#### Meta-analysis archive (ExGrade_v1.0_meta_analysis.zip; 466 files)

results/meta contains, for each of the 92 analysed units (units with at least three clusters; in four metabolome units no feature was measured in three clusters), four files: <UNIT> genes.tsv.gz, the meta-analysis of every feature in the unit (Table 4), including features measured in fewer than three clusters (class Insufficient); <UNIT> genes_G.tsv.gz and <UNIT> genes_V.tsv.gz, matrices of the cluster-level effect sizes and variances (features × study clusters) that entered the meta-analysis; and <UNIT> clusters.tsv, the contributing study clusters with their numbers of contrasts and samples, species and accessions. For 66 units (65 transcriptomic units and the proteome unit), <UNIT> pathways.tsv.gz gives the corresponding pathway results. unit_summary_<LAYER>.tsv lists all units of a layer, including those with fewer than three clusters (status insufficient_unit), with numbers of clusters, contrasts and samples, species composition and counts of features in each class, and inconsistency_calibration.tsv gives the observed and expected shares of inconsistent features for each unit. results/validation contains external_validation_features.tsv.gz (for every feature, unit and atlas: discovery class, pooled *g*, *k*, validation estimate, validation *P*, same sign and replicated), external_validation_summary.tsv (replication rate, sign concordance and Spearman correlation per class, and the Cochran–Armitage trend statistic per unit and atlas), split_half_replication.tsv (replication rate and sign concordance per class, unit and split) and the MoTrPAC effects used for validation, derived from MotrpacRatTraining6moData (CC BY 4.0) by combining sexes (motrpac_8w_effects_METAB_TRNSCRPT.tsv.gz and motrpac_8w_effects_PROT.tsv.gz: feature, log fold change, standard error, number of sexes combined, *z*, *P*, assay, MoTrPAC tissue and major tissue class). results/secondary contains the 18 files (16 tables and 2 JSON files) of the additional analyses described above, including evidence_status_units.tsv (unit status with the quantities used by the status rules), contrast_lambda.tsv (λ, sample size, species, platform and design for the 592 contrasts of ten major transcriptomic units), sensitivity_4_skeletal_muscle.tsv and sensitivity_7_heart.tsv (sensitivity analyses of four skeletal muscle units and of seven units in other tissues, respectively) and reporting_completeness.json. results/numbers.json collects all summary numbers computed from these files.

#### Website archive (ExGrade_v1.0_portal.zip; 135 files)

portal/index.html is the complete single-page application; portal/data contains the unit definitions, status map and checklist (units.json), the search index (index.json), gene synonyms with mouse and rat symbols and Ensembl identifiers (aliases.json), the interface text (i18n.json) and per-feature results in 128 shards (data/s/NNN.json).

#### Code archive (ExGrade_v1.0_code.zip; 67 files)

code/ contains the analysis scripts, numbered in run order (01–06, search and curation; 07, 08 and 10–13, re-processing, effect sizes and validation data; 09, meta-analysis and grading; 14, validation; 15–41, secondary analyses, figures, tables and website data), shared modules and run_downstream.sh (see Usage Notes); protocol/PROTOCOL.md is the analysis protocol with its deviations log; requirements.txt lists the software versions; LICENSE gives the MIT License.

#### Source data

The accessions of all source datasets — 525 GEO series, 16 PRIDE projects and 15 Metabolomics Workbench studies — are given in the curation archive (geo_series.tsv, stage2_series_decisions.tsv and the contrast tables) and, with titles and associated PubMed identifiers, in Supplementary Table 1.

### Data Overview

Table 5 summarizes the re-analysed contrasts by molecular layer, and Fig. 2 shows their distribution over tissues, paradigms and layers. The transcriptome accounts for 800 of the 1,037 re-analysed contrasts (487 RNA-seq and 313 array contrasts) and 365 of the 448 study clusters; skeletal muscle accounts for 52% of transcriptomic study clusters, and 14 of the 21 tissue classes shown in Fig. 2a (including ‘other’, and skin, whose only contrast could not be re-analysed) never reach ten clusters in any combination of paradigm and layer. Of the 107 combinations of tissue class, molecular layer and paradigm shown, 36 have at least three re-analysed independent clusters with shared features and could therefore be meta-analysed, 66 have fewer, and 5 contain only chromatin contrasts, which were catalogued but not re-analysed (Fig. 2a). Females were included in 31% of the 454 GEO discovery study clusters and sex was not reported in 30% (Fig. 2c).

**Table 5.**
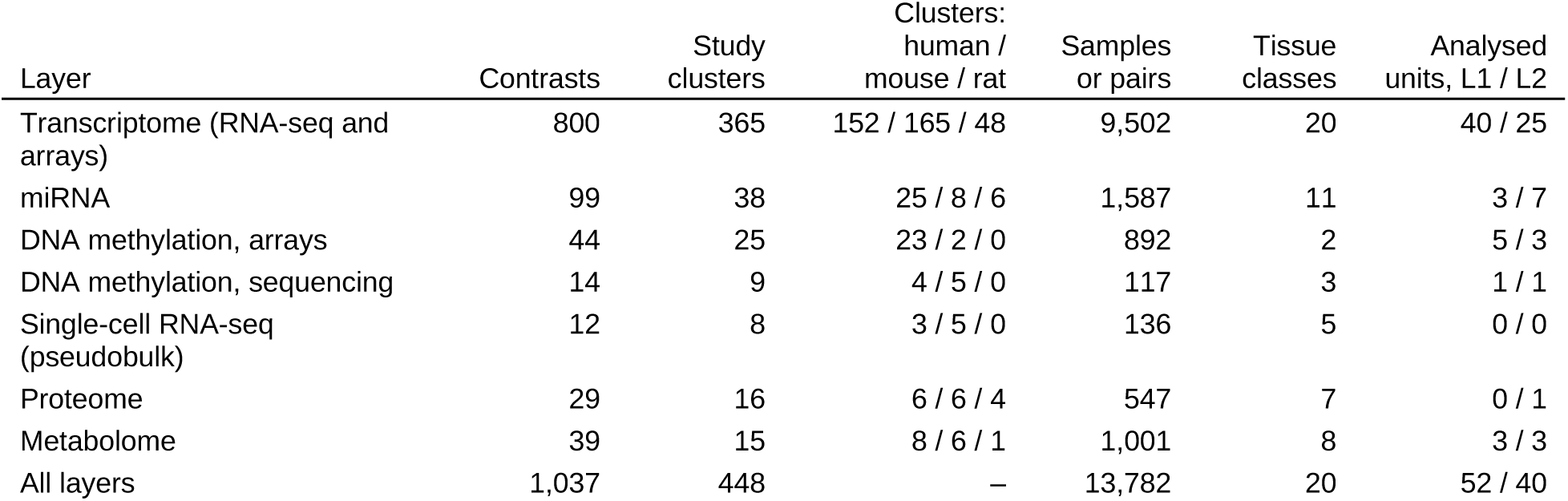
Re-analysed discovery contrasts by molecular layer.

**Fig. 2.**
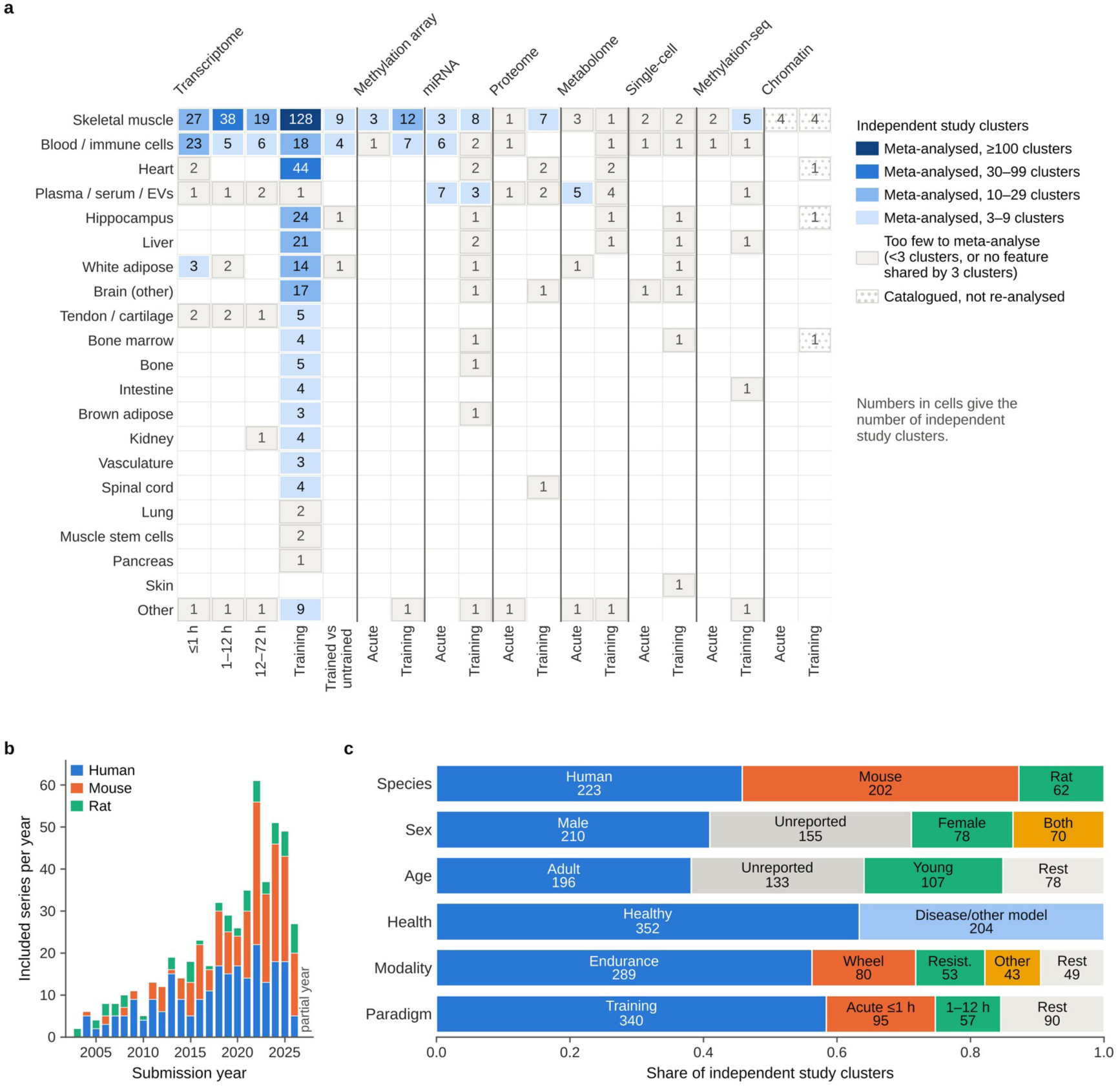
Coverage of the dataset. **a** Number of independent study clusters for each tissue class (rows) by molecular layer and paradigm (columns). Shading indicates whether the combination was meta-analysed (at least three re-analysed clusters with features measured in at least three of them) and, if so, the number of clusters. Dotted cells contain only chromatin contrasts, which were catalogued but not re-analysed; grey cells also include combinations whose contrasts could not be re-analysed, with the number of catalogued clusters shown. For layers other than the transcriptome, the acute time bins are combined into one column, which shows the number of clusters of the most informative time bin; cross-sectional comparisons in these layers, and the spatial and ‘other’ layers, are not shown. **b** Included GEO series by submission year and species; the final year is incomplete. **c** Composition of the discovery study clusters by species, sex, age group, health context, exercise modality and paradigm; clusters contributing contrasts to more than one category are counted in each, and categories with less than 7.5% of clusters are combined as ‘Rest’.

Samples or pairs, exercise plus control samples (independent designs) or matched pairs (paired designs), summed over contrasts. A study cluster profiled in several layers is counted once in the total, and a cluster can include more than one species. Tissue classes include the residual class ‘other’. Analysed units have at least three independent study clusters; L1, modality-specific; L2, all modalities pooled.

### Technical Validation

#### Contrast definitions

All 1,309 GEO contrasts, 29 PRIDE contrasts and 39 Metabolomics Workbench contrasts were resolved programmatically against the repository records: every group contained at least two samples, no sample occurred in both groups of a contrast, and the numbers of selected samples matched the group sizes expected by the curator. All 440 paired discovery contrasts had subject identifiers that matched exercise and control samples. Of the 1,088 GEO discovery contrasts, 969 (89%) were re-analysed; 26 chromatin (accessibility and ChIP-seq), spatial and other contrasts were catalogued but not re-analysed, and 93 could not be re-analysed, most often because only group-level tables, R objects, oversized or unprocessed raw files or targeted arrays had been deposited (12 categories, recorded in unprocessed_reasons.tsv). Of the 487 re-analysed RNA-seq contrasts, 314 used uniformly processed counts (ARCHS4 or NCBI GEO) and 173 used submitter tables, 134 of which required curator-defined column-to-sample maps; each map was checked for the presence and numeric content of all mapped columns and for the expected sample counts.

#### Effect-size calibration

Sample sizes were small: the median transcriptomic contrast had five samples or pairs in its smaller group (interquartile range 3–8; Fig. 3a). To flag contrasts whose effect sizes were shifted across the whole genome, which can result from batch effects aligned with group^36^, from differences in sampling time or from genuinely broad responses, we computed the genomic inflation factor λ of the 592 contrasts in ten major transcriptomic units (Fig. 3b). Contrasts with λ > 2 made up 16–30% of contrasts in the major training units and 33–39% in the acute skeletal muscle units, in which broad transcriptional responses are expected. λ is therefore provided as a diagnostic rather than used as a filter; sensitivity analyses excluding contrasts with λ > 2 are summarized for 11 major units in the sensitivity tables.

**Fig. 3.**
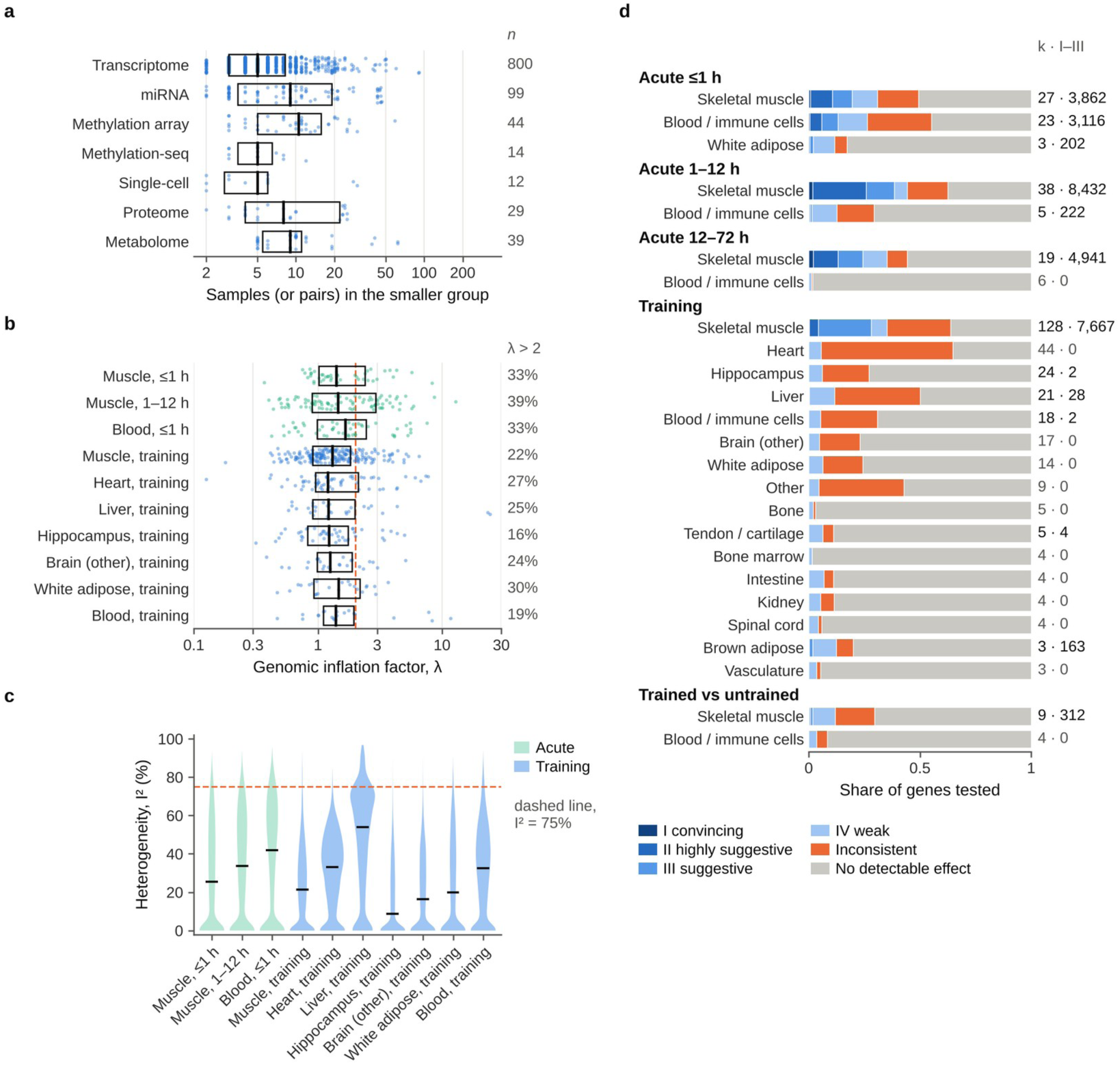
Quality indicators of contrasts and analysis units. **a** Number of samples (or matched pairs, for paired designs) in the smaller group of each re-analysed discovery contrast, by molecular layer; boxes show medians and interquartile ranges, and points show individual contrasts. The number of contrasts is given on the right. **b** Genomic inflation factor λ (median of *g*^2^/var(*g*) across genes divided by 0.4549) of the contrasts in ten major transcriptomic units, displayed as in **a**; the dashed line marks λ = 2, and the share of contrasts with λ > 2 is given on the right. **c** Between-study heterogeneity (*I*^2^) of all genes measured in at least three clusters in the same units; horizontal lines show medians and the dashed line marks *I*^2^ = 75%. **d** Distribution of evidence classes among genes measured in at least three clusters in each level-2 transcriptomic unit (all modalities pooled), grouped by paradigm and ordered by the number of independent clusters (*k*); numbers give *k* and the number of genes in classes I–III.

#### Heterogeneity

The distribution of between-study heterogeneity differed between units (Fig. 3c), and the share of features in each evidence class differed accordingly (Fig. 3d). Because the primary rule for the inconsistent class becomes more likely by chance as the number of clusters increases, the data record gives for each unit the share of features expected to meet this rule under homogeneity and the share meeting a calibrated definition that additionally requires a significant Cochran’s *Q*.

#### Positive controls

Genes with well-established responses to exercise in skeletal muscle^37–39^ were recovered in the expected direction and time course (Fig. 4a). Within 1 h of an acute bout, *NR4A3* was induced (pooled *g* = 3.6; significantly induced in 23 of 24 study clusters and repressed in none), together with *MAFF*, *ATF3*, *EGR1* and *FOS*. At 1–12 h, *PPARGC1A* (*g* = 3.1; significantly induced in 33 of 37 clusters), *PDK4* (*g* = 1.5), *HK2* (*g* = 1.4) and *VEGFA* were induced. After training, transcripts of the mitochondrial enzymes citrate synthase (*CS*; *g* = 0.35, *k* = 124) and *COX4I1* (*g* = 0.40, *k* = 126) were increased (class II), consistent with targeted studies of trained muscle^39^, and *MSTN* was repressed (class III).

**Fig. 4.**
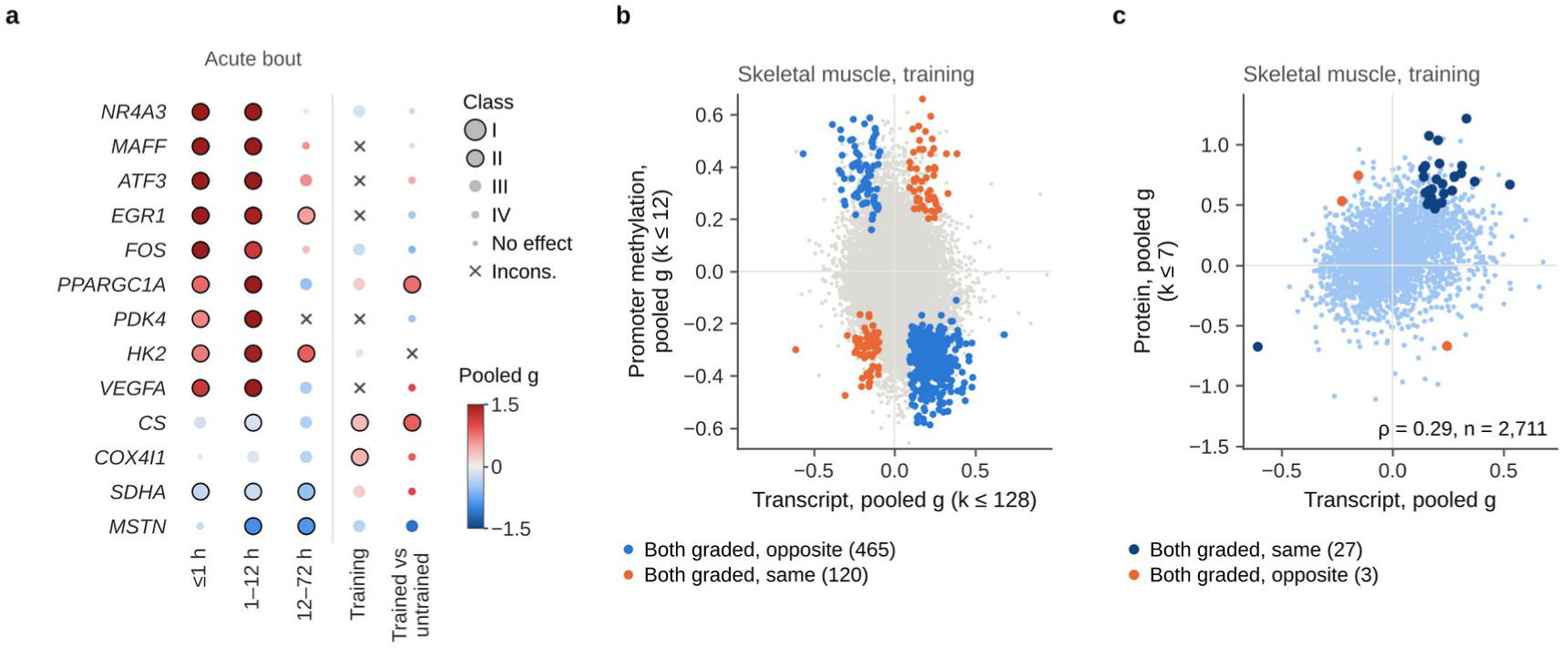
Positive controls and agreement between molecular layers. **a** Pooled effects in skeletal muscle of genes with established responses to exercise, after an acute bout (by time from the end of exercise to sampling), after training and in trained versus untrained people (level-2 units). Colour shows the pooled Hedges’ *g* and symbol size and outline the evidence class (smallest dots, no detectable effect); ×, inconsistent; blank, measured in fewer than three clusters. **b** Pooled effects on promoter DNA methylation (*k* up to 12 clusters) against transcript effects (*k* up to 128) in skeletal muscle after training, for genes measured in at least three clusters in both layers; coloured points are genes graded I–III in both layers that changed in opposite directions (blue, the expected relation) or in the same direction (orange). **c** Pooled effects on protein abundance (*k* up to 7) against transcript effects in skeletal muscle after training; dark blue and orange points are genes graded I–III in both layers that changed in the same direction (the expected relation) or in opposite directions, respectively; ρ, Spearman correlation across all genes shown.

#### Agreement between molecular layers

In the skeletal muscle training unit, which has the most studies in several layers, the layers agreed in the directions expected from their biology, which supports the consistency of the identifier mapping and sign conventions across pipelines. Changes in promoter DNA methylation (array data) were inversely related to transcript changes (Spearman’s ρ = −0.13 across all 16,245 genes measured in both layers and −0.23 among the 5,967 genes graded I–III in the transcriptome), and genes graded I–III in both layers changed in opposite directions nearly four times as often as in the same direction (465 versus 120; Fig. 4b), consistent with the inverse relation between methylation and expression changes in trained human muscle^40^. Protein and transcript effects were positively correlated (ρ = 0.29 across all 2,711 genes and 0.36 among the 1,564 genes graded I–III in the transcriptome), and 27 of the 30 features graded I–III in both layers changed in the same direction (Fig. 4c). Other comparisons, which involve fewer studies, showed weak or no correlation; for example, ρ = 0.00 for sequencing-based promoter methylation in trained muscle and ρ = 0.01 for array methylation within 1 h of exercise (all comparisons are in cross_omic.tsv).

#### Robustness of pooled estimates

Class I requires every leave-one-cluster-out estimate to remain nominally significant in the same direction, and class II requires this of at least 80% of them. In the sensitivity analyses of the skeletal muscle training unit, 79–94% of the 7,667 genes graded I–III remained nominally significant in the same direction when the analysis was restricted to healthy contexts, human studies, RNA-seq or paired designs, and more than 99% after exclusion of the largest cluster, with at least 98% sign concordance in these five analyses. Retention was lower when the analysis was restricted to arrays (74%; *k* = 52), independent-group designs (53%; *k* = 83) or rodent studies (26%; *k* = 71; sign concordance 81%). Results for all 11 major units are provided in the data record.

#### Replication in held-out atlases

In the skeletal muscle training unit, the proportion of genes that replicated in the two held-out atlases (same sign, *P* < 0.05) declined monotonically with evidence class (Fig. 5a,b). In the mouse wheel-running atlas, 79% of class I–II genes (*n* = 611) replicated, compared with 58% of class III, 42% of class IV and 16% of genes without a detectable effect (Cochran–Armitage *z* = 23.3). In the MoTrPAC rat treadmill-training atlas, the corresponding figures were 37% (*n* = 618), 19%, 14% and 8% (*z* = 11.7). Every modality-specific muscle training unit with at least three classes tested showed a positive trend in both atlases (*z* = 15.0–34.6 in mouse and 9.5–22.5 in rat; Fig. 5c), and pooled effects of class I–III genes in the wheel-running unit correlated with the mouse wheel-running atlas (ρ = 0.80, *n* = 1,323; Fig. 5e). In MoTrPAC’s muscle proteome, 8 of the 9 class I–II proteins with rat orthologues replicated, compared with 45% of class III, 37% of class IV and 23% of proteins without a detectable effect (*z* = 7.1). Outside skeletal muscle, few features reached classes I–III, so trend tests rested mainly on the weak and null classes and gave mixed results (*z* from −3.1 to 11.5); the 163 genes graded I–III in brown adipose tissue after training, a unit with three clusters, did not replicate in either atlas (0% and 5%, compared with 2% and 11% of genes without a detectable effect). The atlases cover training only, so the acute and cross-sectional classes could not be tested externally.

**Fig. 5.**
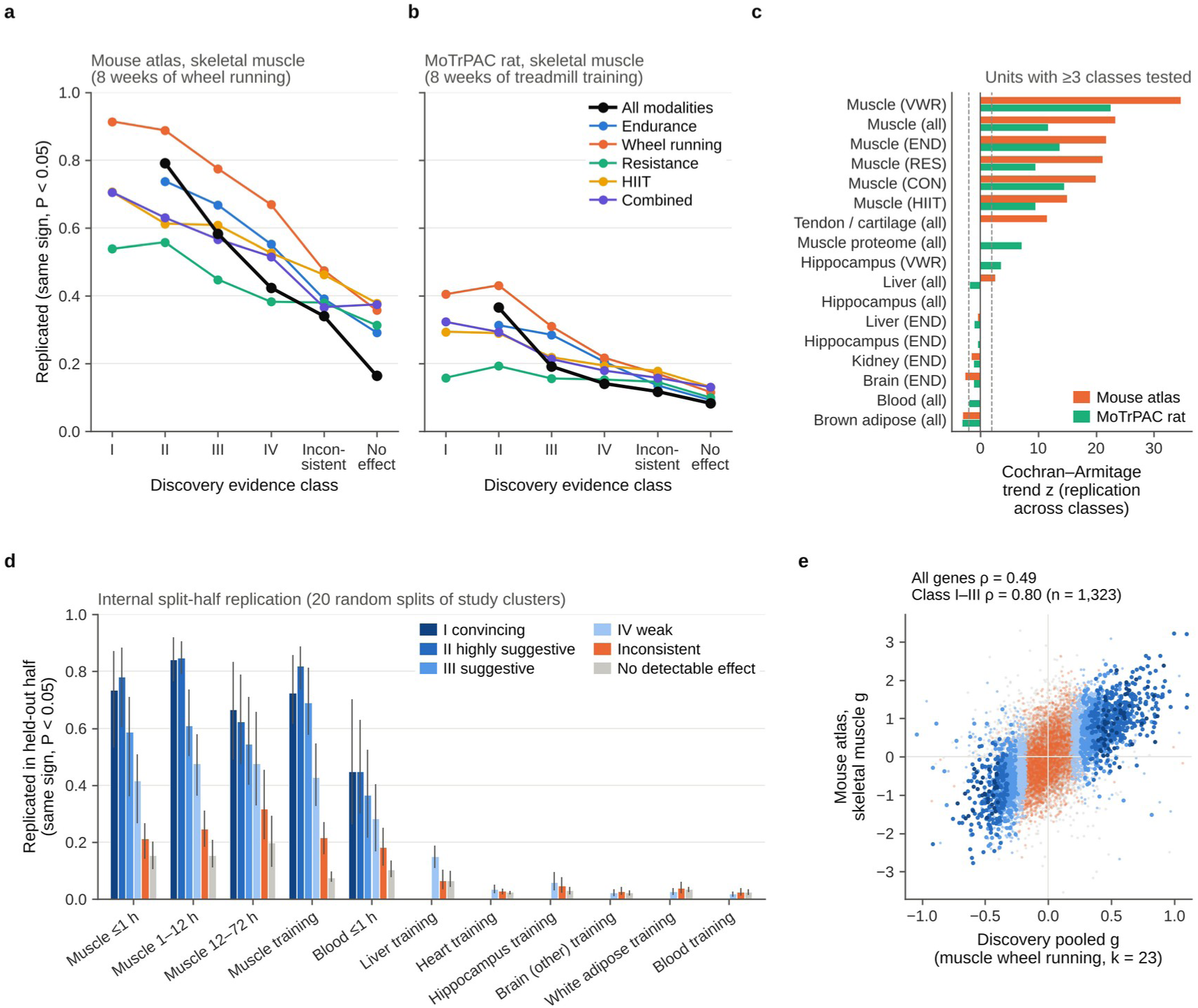
Replication of evidence classes in held-out data. **a**, **b** Proportion of skeletal muscle training genes that replicated, by discovery evidence class, in the held-out mouse atlas of Maier et al. (8 weeks of voluntary wheel running; **a**) and in MoTrPAC rats (8 weeks of treadmill training; **b**). Replication is defined as an effect of the same sign with *P* < 0.05 in the validation data; lines show the level-2 unit (all modalities) and each modality-specific unit, and classes with fewer than 20 genes are omitted. **c** Cochran–Armitage trend statistics for replication across classes (I, II, III, IV and no detectable effect) for every unit with at least three classes tested, including the skeletal muscle proteome (MoTrPAC only); dashed lines, |*z*| = 1.96. **d** Internal split-half replication in level-2 transcriptomic units over 20 random splits of the study clusters into halves; bars show means over the splits in which a class contained at least ten genes, and lines the 10th–90th percentiles. **e** Pooled effects in the discovery skeletal muscle wheel-running unit against effects in the mouse atlas, coloured by evidence class; axes are truncated at the 99.8th percentile of absolute effects (52 of 13,164 genes lie outside and are not shown); genes measured in fewer than three discovery clusters are not shown but are included in the correlation for all genes. END, endurance; VWR, voluntary wheel running; RES, resistance; HIIT, high-intensity interval training; CON, combined.

#### Split-half replication

When the study clusters of each transcriptomic unit with at least eight clusters were split at random into halves 20 times, class I–II genes graded in one half replicated in the other, on average over the splits in which a class contained at least ten genes, in 73–85% of cases in skeletal muscle within 12 h of exercise, 72–82% in the muscle training unit and 45% in blood within 1 h of exercise, against 8–15% for genes without a detectable effect (Fig. 5d). In the training units of heart, hippocampus, other brain regions, white adipose tissue and blood, half-samples yielded no class I–II genes, and the weak, inconsistent and null classes replicated at 2–6%, close to the 2.5% expected by chance.

#### Reproducibility

To check that the deposited files are complete and self-consistent, we unpacked the code, curation and effect-size archives into an empty folder and re-ran the meta-analysis and grading script (code/09_meta_grade.py) for all seven layers, followed by the inconsistency calibration (code/20_inconsistency.py), changing only the hard-coded base paths in the two scripts. All 442 files of results/meta — the feature-level meta-analyses, cluster-level matrices and cluster lists of the 92 analysed units, the pathway results, the unit summaries and the inconsistency calibration — were identical to the deposited files after decompression.

### Usage Notes

#### Choosing units and interpreting classes

L2 units pool exercise modalities and L1 units are modality-specific; the time bins (A0, A1, A2, C and X) separate acute responses by sampling time from training and cross-sectional comparisons. Evidence classes rate the credibility of an average effect across heterogeneous designs; they do not measure its magnitude or biological importance, and the sign of the pooled *g* gives the direction. Classes from units with three to five clusters should be treated as provisional, because a Wald test with a restricted-maximum-likelihood estimate of τ^2^ can be anticonservative when there are few clusters. External tests were possible mainly for skeletal muscle training units; the classes of other tissues have not been tested in independent data. In large units, class I can be dominated by features measured in fewer, more homogeneous studies: in the skeletal muscle training unit, class I genes were measured in a median of 18 clusters, against 96 for class II, and 29% of them are protein-coding. Users who select reference sets should therefore also consider *k* and gene type. Classes will change as studies accumulate.

#### Re-using effect sizes

The per-contrast files can be joined to the contrast annotations through the contrast identifier (key = gse + ’ ’ + cid). New pooled estimates for alternative groupings can be computed with standard meta-analysis software from the cluster-level matrices (<UNIT> genes_G.tsv.gz and <UNIT> genes_V.tsv.gz) or, for groupings that cross units, from the per-contrast files after combining contrasts within study clusters as described in the Methods; code/09_meta_grade.py implements these steps. Effect sizes for paired and independent designs are on the same standardized scale (*d*_av_ for paired designs). Rodent-specific genes are not represented, and features from different layers share identifiers only through gene-level mapping.

#### Annotations and their limits

Contrast annotations were derived from the repository metadata, so they are only as complete as the deposited records: sex was not reported for 30% of discovery study clusters and age group for 28%; the time from the end of exercise to sampling was missing for 61 of 445 acute contrasts; and the time since the last bout was recorded for only 64 of 612 training contrasts. Annotation was performed by a large language model under written rules and verified programmatically for sample selection only; the exclusion decisions and the other annotations (tissue, timing, modality, health context and sex) were not audited against a manual reference, so misclassification of such details remains possible. Contrasts with genomic inflation in the major transcriptomic units can be identified in contrast_lambda.tsv. The search covered GEO, PRIDE and Metabolomics Workbench only, so controlled-access data and studies deposited elsewhere are not included, and SubSeries of matched GEO SuperSeries that did not themselves match the search terms were not screened.

#### Licences of the source data

GEO and PRIDE data were used under the terms of these repositories, and the MoTrPAC data package is distributed under CC BY 4.0. The Metabolomics Workbench studies used here are distributed under the CC BY-NC-ND 4.0 licence; the data record contains no source measurements from them, only per-metabolite summary statistics (in the metabolome effect-size, meta-analysis and validation files), and users who reuse these files should take the source licence into account. Users who return to the source data should follow the licence and citation requirements of each source.

#### Running the website and the code

The website loads its data with fetch requests, so it must be served over HTTP rather than opened as a local file: after unpacking ExGrade_v1.0_portal.zip, run python3 -m http.server 8000 in the portal folder and open http://localhost:8000. The meta-analysis, grading and inconsistency calibration can be reproduced from the deposited files: unpack the code, curation and effect-size archives into one folder, install Python 3.13 and the packages in requirements.txt, set the base paths at the top of code/09_meta_grade.py and code/20_inconsistency.py to that folder and run the two scripts (see Technical Validation). The other scripts document the complete pipeline but cannot be re-run from the record alone: the re-processing scripts (01–08 and 10–13) download source data from GEO, ARCHS4, PRIDE, Metabolomics Workbench and MoTrPAC and additionally need fsspec, aiohttp and py7zr; these and several downstream scripts read reference files (NCBI gene annotations and 1:1 orthologue tables, GEO platform annotations, gene-set files and promoter coordinates) that are obtained from public sources but are not part of the record; and code/run_downstream.sh, which documents the order in which the downstream scripts were run, starts with the proteome and metabolome effect sizes (script 13), which require the downloaded source tables. Re-processing with later ARCHS4 or Reactome releases may give slightly different results.

## Data Availability

The data record is available at Zenodo (https://doi.org/10.5281/zenodo.23642143)^35^ under a CC BY 4.0 licence. The source data are available from GEO, PRIDE and Metabolomics Workbench under the accessions listed in the data record and in Supplementary Table 1. The held-out validation data are available from the MoTrPAC data package^33^ and from GEO (GSE344069)^34^.

## Code Availability

The code used for searching, curation, re-processing, meta-analysis, grading, validation, secondary analyses and the query website is included in the data record (ExGrade_v1.0_code.zip)^35^ under the MIT License, together with the code for Fig. 5 (code/28_fig6_validation.py). The scripts for Figs. 1–4 and the summary files they read are provided as Supplementary Software. The analyses were run with Python 3.13.17, NumPy 2.5.3, SciPy 1.18.1, pandas 3.0.5, h5py 3.16.0, pyreadr 0.5.7 and matplotlib 3.11.2; the download steps also used fsspec, aiohttp and py7zr. Parameters are set in the scripts and described in the Methods, and the reference files that some scripts need are described in the Usage Notes.

## Supporting information

Supplementary Software

## References

1. Pedersen, B.K. & Saltin, B. Exercise as medicine – evidence for prescribing exercise as therapy in 26 different chronic diseases. Scand. J. Med. Sci. Sports 25 (Suppl. 3), 1–72 (2015).

2. Neufer, P.D. et al. Understanding the cellular and molecular mechanisms of physical activity-induced health benefits. Cell Metab. 22, 4–11 (2015).

3. Hawley, J.A., Hargreaves, M., Joyner, M.J. & Zierath, J.R. Integrative biology of exercise. Cell 159, 738–749 (2014).

4. Button, K.S. et al. Power failure: why small sample size undermines the reliability of neuroscience. Nat. Rev. Neurosci. 14, 365–376 (2013).

5. Ioannidis, J.P. Why most published research findings are false. PLoS Med. 2, e124 (2005).

6. Pillon, N.J. et al. Transcriptomic profiling of skeletal muscle adaptations to exercise and inactivity. Nat. Commun. 11, 470 (2020).

7. Amar, D. et al. Time trajectories in the transcriptomic response to exercise – a meta-analysis. Nat. Commun. 12, 3471 (2021).

8. Jacques, M. et al. Molecular landscape of sex- and modality-specific exercise adaptation in human skeletal muscle through large-scale multi-omics integration. Cell Rep. 44, 115750 (2025).

9. Voisin, S. et al. Exercise is associated with younger methylome and transcriptome profiles in human skeletal muscle. Aging Cell 23, e13859 (2024).

10. Pan, L. et al. ExerGeneDB: A physical exercise-regulated differential gene expression database. J. Sport Health Sci. 14, 101027 (2025).

11. Sun, L. et al. GEPREP: A comprehensive data atlas of RNA-seq-based gene expression profiles of exercise responses. J. Sport Health Sci. 14, 100992 (2025).

12. Li, Z., Xu, Z. & Li, X. SportsXbiodata: A web platform for integrative analysis of exercise-induced gene and protein expression. iScience 29, 114403 (2026).

13. Sanford, J.A. et al. Molecular Transducers of Physical Activity Consortium (MoTrPAC): mapping the dynamic responses to exercise. Cell 181, 1464–1474 (2020).

14. MoTrPAC Study Group. Temporal dynamics of the multi-omic response to endurance exercise training. Nature 629, 174–183 (2024).

15. Maier, G. et al. Multi-omics atlas of chronic exercise. Cell 10.1016/j.cell.2026.09.044 (2026).

16. Ioannidis, J.P. Integration of evidence from multiple meta-analyses: a primer on umbrella reviews, treatment networks and multiple treatments meta-analyses. CMAJ 181, 488–493 (2009).

17. Fusar-Poli, P. & Radua, J. Ten simple rules for conducting umbrella reviews. Evid. Based Ment. Health 21, 95–100 (2018).

18. Page, M.J., et al. The PRISMA 2020 statement: an updated guideline for reporting systematic reviews. BMJ 372, n71 (2021).

19. Barrett, T. et al. NCBI GEO: archive for functional genomics data sets—update. Nucleic Acids Res. 41, D991–D995 (2013).

20. Perez-Riverol, Y. et al. The PRIDE database resources in 2022: a hub for mass spectrometry-based proteomics evidences. Nucleic Acids Res. 50, D543–D552 (2022).

21. Sud, M. et al. Metabolomics Workbench: An international repository for metabolomics data and metadata, metabolite standards, protocols, tutorials and training, and analysis tools. Nucleic Acids Res. 44, D463–D470 (2016).

22. Lachmann, A. et al. Massive mining of publicly available RNA-seq data from human and mouse. Nat. Commun. 9, 1366 (2018).

23. Robinson, M.D. & Oshlack, A. A scaling normalization method for differential expression analysis of RNA-seq data. Genome Biol. 11, R25 (2010).

24. Smyth, G.K. Linear models and empirical bayes methods for assessing differential expression in microarray experiments. Stat. Appl. Genet. Mol. Biol. 3, Article3 (2004).

25. Marot, G., Foulley, J.L., Mayer, C.D. & Jaffrézic, F. Moderated effect size and P-value combinations for microarray meta-analyses. Bioinformatics 25, 2692–2699 (2009).

26. Morris, S.B. & DeShon, R.P. Combining effect size estimates in meta-analysis with repeated measures and independent-groups designs. Psychol. Methods 7, 105–125 (2002).

27. Liberzon, A. et al. The Molecular Signatures Database (MSigDB) hallmark gene set collection. Cell Syst. 1, 417–425 (2015).

28. Milacic, M. et al. The Reactome Pathway Knowledgebase 2024. Nucleic Acids Res. 52, D672–D678 (2024).

29. Viechtbauer, W. Bias and efficiency of meta-analytic variance estimators in the random-effects model. J. Educ. Behav. Stat. 30, 261–293 (2005).

30. Higgins, J.P. & Thompson, S.G. Quantifying heterogeneity in a meta-analysis. Stat. Med. 21, 1539–1558 (2002).

31. IntHout, J., Ioannidis, J.P., Rovers, M.M. & Goeman, J.J. Plea for routinely presenting prediction intervals in meta-analysis. BMJ Open 6, e010247 (2016).

32. Egger, M., Davey Smith, G., Schneider, M. & Minder, C. Bias in meta-analysis detected by a simple, graphical test. BMJ 315, 629–634 (1997).

33. Gay, N., Jean Beltran, P., Amar, D., Jimenez-Morales, D. & MoTrPAC Study Group. MotrpacRatTraining6moData: data for analysis of the MoTrPAC endurance exercise training study in 6-month-old rats, version 2.0.0. *GitHub* https://github.com/MoTrPAC/MotrpacRatTraining6moData (accessed October 2026).

34. Maier, G., et al. Multi-Omics Atlas of Chronic Exercise [ChronicRW_RNAseq]. Gene Expression Omnibus https://identifiers.org/geo:GSE344069 (2026).

35. Li, C. ExGrade v1.0: code and data for “Molecular effects of exercise training replicate mainly in skeletal muscle”. Zenodo 10.5281/zenodo.23642143 (2026).

36. Leek, J.T. et al. Tackling the widespread and critical impact of batch effects in high-throughput data. Nat. Rev. Genet. 11, 733–739 (2010).

37. Egan, B. & Sharples, A.P. Molecular responses to acute exercise and their relevance for adaptations in skeletal muscle to exercise training. Physiol. Rev. 103, 2057–2170 (2023).

38. Pilegaard, H., Saltin, B. & Neufer, P.D. Exercise induces transient transcriptional activation of the PGC-1α gene in human skeletal muscle. J. Physiol. 546, 851–858 (2003).

39. Perry, C.G. et al. Repeated transient mRNA bursts precede increases in transcriptional and mitochondrial proteins during training in human skeletal muscle. J. Physiol. 588, 4795–4810 (2010).

40. Lindholm, M.E. et al. An integrative analysis reveals coordinated reprogramming of the epigenome and the transcriptome in human skeletal muscle after training. Epigenetics 9, 1557–1569 (2014).

